# Differential subcortical–cortical circuit reorganization in autism and ADHD

**DOI:** 10.64898/2026.10.13.758196

**Authors:** Guan Zhou, Ke Xie, Alexander Ngo, Rui Ding, Donna Gift Cabalo, Yezhou Wang, Meghan Smith, Misha Nakamura, Jessica Royer, Naomi Serediuc, Paule J. Toussaint, Laurent Mottron, Adriana Di Martino, Michael P. Milham, Seok-Jun Hong, Alan C. Evans, Boris C. Bernhardt

## Abstract

Autism spectrum disorder (ASD) and attention-deficit/hyperactivity disorder (ADHD) often co-occur and share symptoms as well as genetic risk factors. While subcortical-cortical networks have been implicated in both conditions, particularly involving thalamic and striatal circuits, patterns of shared and diverging disruptions remain unclear. Here we integrated resting-state functional connectivity (FC) and regression dynamic causal modeling (rDCM) across a transdiagnostic multi-site cohort. We used statistical group comparisons and individualized normative deviation mapping to map network deviations in ASD and ADHD relative to neurotypical controls and against one another. Multivariate associative techniques examined associations to clinical and cognitive phenotypes. Our transdiagnostic analyses revealed significant differentiation of ASD and ADHD across subcortical-cortical circuits. While ASD was characterized by widespread thalamocortical hyperconnectivity and bidirectional elevated coupling across thalamic subdivisions, ADHD exhibited a focal pattern of striatocortical disruption, anchored predominantly within the caudate. Multivariate analyses identified shared and disorder-specific brain-behavior associations. Across both conditions, aberrant striatocortical signaling was associated with reduced overall intelligence measures. Clinically, ASD-related thalamocortical connectivity disruptions mapped onto generalized cognitive impairment, whereas ADHD-related subcortical alterations tracked hyperactive and inattentive symptom dimensions. Our findings differentiate ASD and ADHD through distinct patterns of functional and directional subcortical-cortical dysregulations.

## Introduction

ADHD and ASD are common and generally lifelong neurodevelopmental conditions. Despite their distinct symptoms (Castellanos et al., 2006; Lai et al., 2019; Rommelse et al., 2010), both conditions frequently co-occur and have been related to common risk factors (Lee et al., 2019; Mattheisen et al., 2022; Verhoef et al., 2021) as well as partially shared cognitive profiles, particularly atypical executive functioning and attentional control (Craig et al., 2016; Schachar et al., 2023). This overlap has challenged current diagnostic boundaries and highlighted the need to identify shared as well as disorder-specific neurobiological substrates (Astle et al., 2022). However, an increasing body of evidence cautions against over-emphasizing shared mechanisms, demonstrating robust neurobiological, cognitive, and developmental divergence between the two conditions (Bedford et al., 2025; Lau-Zhu et al., 2019; Watanabe & Watanabe, 2023). Given these conflicting perspectives, there is a critical need to implement analytical frameworks capable of simultaneously disentangling shared from disorder-specific neurobiological substrates.

A growing body of evidence conceptualizes both ADHD and ASD as “connectopathies” *i.e.* disruptions of large-scale brain networks, with disruptions observed in cortico-subcortical circuits, particularly those involving the thalamus and the striatum (Di Martino et al., 2014; Di Martino et al., 2013; Faraone & Larsson, 2019). Neuroanatomically, these subcortical hubs play specialized and complementary roles in regulating cognitive and behavioral output. The thalamus acts as a key modulator of whole brain dynamics (Jones, 2001; Müller et al., 2020; Oldham & Ball, 2023; Shine et al., 2023), regulating sensorimotor as well as higher order cognitive processing (Halassa & Kastner, 2017). In contrast, the striatum, comprising the caudate and putamen, is embedded in recurrent cortical-subcortical loops characterized by their topographic organization and intrinsic hierarchical dynamics, facilitating the integration of parallel cortical input (Choi et al., 2012; Haber, 2016). Across typical neurodevelopment, subcortico-cortical networks undergo marked functional refinement, shifting from more diffuse connectivity in childhood toward increasingly segregated and specialized thalamocortical and frontostriatal circuits in adulthood (Greene et al., 2020; Supekar et al., 2009). Consequently, structural and functional alterations within subcortical– cortical systems have been increasingly implicated in the pathophysiology of both conditions. In ASD, thalamocortical hyper-connectivity has been suggested to reflect to sensory over-responsiveness and gating deficits (Green et al., 2017; Nair et al., 2015; Uddin et al., 2013), whereas ADHD is prominently characterized by frontostriatal and thalamocortical hypo-connectivity associated with executive dysfunction and inhibitory deficits (Castellanos & Proal, 2012; Cortese et al., 2012; Mills et al., 2012).

Resting-state functional magnetic resonance imaging (rs-fMRI) has emerged as a key technique for studying intrinsic functional networks across the entire brain with high spatial precision and reliability (Biswal et al., 1995; Fox & Raichle, 2007; Shehzad et al., 2009; Smith et al., 2009). Beyond mapping large-scale functional connectivity (FC) topographies in typical development (Luo et al., 2024; Park et al., 2022; Sydnor et al., 2023), rs-fMRI has firmly established ASD and ADHD as complex network disorders characterized by robust yet heterogeneous patterns of atypical connectivity (Castellanos & Proal, 2012; Cortese et al., 2012; Di Martino et al., 2014; Hull et al., 2017; King et al., 2019; Martino et al., 2011; Menon, 2011; Van Den Heuvel & Sporns, 2019; Zuo et al., 2010). However, traditional FC analyses assess undirected statistical associations between regional BOLD signals. While these static topographical maps identify where networks are disrupted, they lack directionality, making it impossible to distinguish between driving and driven neural influences (Fornito et al., 2015; Friston, 2011). Consequently, it remains unclear whether the observed network hyper-or hypoconnectivity reflects specific signal flow alterations from the cortex to subcortex, or *vice versa*. Recent advances in generative modeling, specifically regression dynamic causal modelling (rDCM), can overcome this limitation by capturing both the direction and strength of inter-regional signaling (Frässle, Aponte, et al., 2021). Prior work has shown to utility of rDCM approaches to study intrinsic flow patterns in cortical systems (Paquola et al., 2019; Paquola et al., 2025), as well as disruptions of flow in neurological (Xie et al., 2024), and neuropsychiatric conditions such as schizophrenia and major depression (Frässle, Harrison, et al., 2021; Stephan et al., 2010). As a result, rDCM stands as a promising technique to clarify the directional nature of FC alterations in ASD and ADHD.

While traditional case-control designs and between-group comparisons have been instrumental in delineating group-average alterations across clinical cohorts, they implicitly assume group-wise homogeneity and may obscure marked inter-individual variability inherent to neurodevelopmental conditions. To complement group-level inferences and better resolve this underlying heterogeneity, research has increasingly incorporated individual-specific analytical frameworks. In this context, normative deviation mapping has emerged as a framework to quantify single-subject deviations relative to expected developmental trajectories (Marquand et al., 2016; Rutherford et al., 2022). By establishing statistical benchmarks of typical variance, akin to pediatric growth charts, these models map individualized profiles of atypical brain organization, acknowledging that the neurobiological instantiation of developmental conditions can vary markedly from person to person (Bethlehem et al., 2022; Sydnor et al., 2023; Wolfers et al., 2018).

In parallel, capturing the complexity of these conditions requires analytical strategies capable of linking individualized neurobiological profiles to clinical and cognitive phenotypes. Multivariate associative frameworks, such as partial least squares (PLS), have been widely adopted for this challenge. They allow identifying distributed patterns of brain structure and function that reliably covary with measures of behavior and clinical severity (McIntosh & Lobaugh, 2004; Mišić et al., 2016; Zeighami et al., 2019). By applying multivariate techniques to individualized connectivity features, recent work has identified complex covariations in clinical cohorts (Kebets et al., 2019; Mihalik et al., 2022; Mišić et al., 2016; Rodríguez-Cruces et al., 2020; Xia et al., 2018), mapping how distributed network dysregulation associates with diverse, overlapping symptom dimensions. To delineate the shared and condition-specific neural features of ASD and ADHD, the present study profiled the functional organization of subcortical-cortical circuits in a large multisite cohort. We applied rs-fMRI connectivity analysis and rDCM to profile thalamo-cortical and striato-cortical network organization and compare these features across clinical and typically developing groups. We complement these results with an individualized normative deviation framework via standardized W-scores to map deviations relative to a typically developing baseline to ensure the robustness of our findings across analytical frameworks. We furthermore used multivariate associative techniques to study associations between these directional network alterations and multidimensional clinical and cognitive profiles. Guided by prior literature implicating divergent cortico-subcortical circuits in these conditions, we anticipated that while ASD and ADHD may exhibit overlapping macroscale connectivity alterations, high-resolution directed modeling and normative mapping would resolve condition-specific divergences in signal flow and their corresponding behavioral associations.

## Methods

### MRI acquisition

High-resolution T1-weighted structural MRI and rs-fMRI data were acquired across 21 international scanning sites aggregated in the ABIDE-1, ADHD-200, and HBN initiatives. Given the large-scale, cross-consortium nature of this study, imaging protocols and scanner platforms varied across datasets. Most data were collected using 3T MRI systems (*e.g.,* Siemens Allegra/TrioTim/Prisma, Philips Achieva/Intera, and GE platforms), with the exception of one HBN site that collected data on a 1.5T Siemens Avanto system. Across all sites, rs-fMRI data were predominantly acquired using gradient-echo Echo Planar Imaging (EPI) sequences, with Repetition Times (TR) ranging from 800 to 3000 ms, tailored to individual site protocols.

A catalog of scanner models and sequence parameters for all 21 sites (15 from ABIDE-1, 2 from ADHD-200, and 4 from HBN) is provided in **Supplementary Table S2**. Descriptions of the original scanning protocols can also be found in the consortium data descriptors (Alexander et al., 2017; consortium, 2012; Di Martino et al., 2014). As outlined below, hardware and sequence heterogeneity was mitigated using ComBat harmonization (Fortin et al., 2017) directly to individual-level connectivity metrics (including static FC matrices and EC parameters) prior to statistical and normative deviation mapping.

### MRI processing

While rs-fMRI preprocessing utilized C-PAC universally (Craddock et al., 2013), cortical surface extraction and volume-to-surface mapping strategies varied slightly to accommodate preprocessing pipelines provided by each consortium.

Processed ADHD-200 data were obtained from the Preprocessed Connectomes Project (http://fcon_1000.projects.nitrc.org/indi/adhd200/). Structural T1-weighted images were preprocessed using FreeSurfer (Dale et al., 1999; Fischl, 2012), encompassing intensity normalization, skull stripping, and tissue segmentation, followed by surface inflation and spherical registration to the fsaverage template. Functional images were co-registered to MNI152 space through linear and nonlinear transformations. rs-fMRI time series were mapped to individual cortical mid-thickness surfaces and resampled to the fsLR 10k template (Glasser et al., 2013). After comprehensive preprocessing and quality assurance, a total of 318 participants remained from two sites in the ADHD-200 dataset: New York University (N = 148; 78 ADHD, 70 TD) and Peking University (N = 170; 66 ADHD, 104 TD).

Data for ABIDE-1 and HBN were accessed via the Preprocessed Connectomes Project (http://preprocessed-connectomes-project.org/abide/) and the FCP/INDI repository (http://fcon_1000.projects.nitrc.org/indi/cmi_healthy_brain_network/), respectively. For both datasets, C-PAC (Craddock et al., 2013) directly integrated CIVET (MacDonald et al., 2000) for structural processing and surface extraction. Similar to FreeSurfer, CIVET performs intensity non-uniformity correction, linear registration to MNI152 space, tissue classification into gray matter, white matter, and cerebrospinal fluid, and extraction of white matter and pial surfaces using deformable models. Functional volumes for these cohorts were directly mapped to the native cortical surfaces at CIVET 20k resolution without an intermediate volumetric MNI co-registration, which minimized interpolation artifacts during mapping. Following our unified quality assurance procedures and harmonization requirements, the final analysis included 652 participants across 13 sites from the ABIDE-1 dataset (N = 652; 303 ASD, 349 TD) and 416 participants across 4 sites from the HBN dataset (N = 416; 23 ASD, 304 ADHD, 89 TD).

Both the cortical surface and volumetric subcortical data were spatially smoothed using a 6 mm full-width-at-half-maximum (FWHM) Gaussian kernel. Cortical surface meshes for the ABIDE and HBN cohorts were downsampled to fsLR 10k space to align with the ADHD-200 cohort, and subcortical structures across all datasets were resampled to align with the Human Connectome Project (HCP) subcortical template space (Glasser et al., 2013).

#### Quality control (QC) and age range

Structural QC involved the evaluation of signal-to-noise ratios as well as visual inspection to exclude participants with major imaging artifacts, topological defects, or failures in cortical surface segmentation. For functional QC, temporal derivative measures and framewise displacement (Power et al., 2012) were computed. Participants exhibiting head motion beyond a mean FD≥0.3 mm were excluded. We furthermore restricted our analysis to school-aged children and adolescents between 7 and 18 years of age. Following these procedures, the final analysis cohort comprised 1,386 participants (325 ASD, 448 ADHD, and 613 typically developing [TD] controls). Demographic and clinical characteristics are summarized in **Table 1**. A comprehensive, site-by-site breakdown of sample sizes, attrition rates, and head motion profiles is detailed in **Supplementary Table S1**.

**Table 1.** Demographic and clinical characteristics of the combined cohort.

| Characteristic | ASD (n=325) | ADHD (n=448) | TD (n=613) | Statistic | p-value |
| --- | --- | --- | --- | --- | --- |
| Age (years) | 12.90 ± 2.84 | 11.94 ± 2.92 | 12.47 ± 2.74 | F = 11.38 | < 0.001 |
| Sex (Male, n (%)) | 277 (85.2%) | 314 (70.1%) | 432 (70.5%) | $\chi^2 = 28.67$ | < 0.001 |
| Mean FD (mm) | 0.11 ± 0.06 | 0.12 ± 0.04 | 0.10 ± 0.06 | F = 29.48 | < 0.001 |
| Full-scale IQ | 105.08 ± 17.57<br>(n=304) | 102.68 ± 16.02<br>(n=387) | 111.28 ± 13.76<br>(n=561) | F = 38.90 | < 0.001 |
| Verbal IQ | 104.09 ± 18.86<br>(n=287) | 106.60 ± 15.74<br>(n=388) | 112.49 ± 14.47<br>(n=540) | F = 30.38 | < 0.001 |
| Performance IQ | 104.92 ± 18.05<br>(n=286) | 101.11 ± 15.61<br>(n=388) | 107.71 ± 14.30<br>(n=538) | F = 20.01 | < 0.001 |

**Notes:** Values are presented as mean ± SD or n (%). **Abbreviations:** ASD, autism spectrum disorder; ADHD, attention-deficit/hyperactivity disorder; TD, typically developing; FD, framewise displacement; IQ, intelligence quotient. Statistical comparisons were performed using one-way ANOVA for continuous variables and Pearson's chi-square test for categorical variables.

Clinical symptom severity was assessed using cohort-specific measures (*e.g.,* ADOS for ABIDE, SRS for HBN). To harmonize metrics for transdiagnostic multivariate analyses, symptom scores were standardized (Z-scored) within each cohort. A summary of the raw clinical symptom profiles across cohorts is provided in **Supplementary Table S3**.

### FC analysis

#### Subregion definition

Consistent with our *a-priori* hypotheses focused on cortico-subcortical connectopathies, and our subcortical regions of interest (ROIs) were restricted to 8 thalamic and 10 striatal subdivisions based on the Melbourne Subcortex Atlas (Tian et al., 2020). This parcellation was selected because it is functional-connectivity-driven, ensuring high intra-regional functional homogeneity that aligns with our resting-state fMRI substrate and avoids artificial signal averaging typical of purely anatomical delineations. Detailed definitions are provided in **Supplementary Table S4**.

*Subdivision dominance analysis.* To quantify the contribution of individual subcortical subdivisions to significant FC alterations, we employed dominance analysis (Azen & Budescu, 2003). This variance decomposition method estimates the relative importance of each subdivision predictor in a multiple regression framework. Crucially, analysis was performed independently within the specific positive and negative FC masks identified in the prior random field theory (RFT) step. The significance of each subdivision’s dominance was established using 5,000 spatial permutations, with p-values stringently corrected using the Benjamini-Hochberg false discovery rate (FDR) procedure (p < 0.05) (Benjamini & Hochberg, 1995).

#### Network contextualization

To contextualize these pathological alterations within established cognitive architectures, we mapped the unthresholded, whole-brain Cohen’s d effect size maps onto the canonical Yeo 7-network parcellation (Yeo et al., 2011).

*Spatial null models*. To validate the robustness of the subcortical-cortical connectivity profiles, we evaluated their spatial consistency across individual scanning sites. For each diagnostic group, we calculated the spatial correlations between the global analytical mean connectivity map and the maps derived from each independent site (n ≥ 5). To establish significance while controlling for the spatial autocorrelation, we generated spatial null models using Moran Spectral Randomization (MSR) with 5,000 iterations (Vos de Wael et al., 2020). MSR generates surrogate maps by randomizing the coefficients of the Laplacian eigenvectors of the cortical graph, thereby preserving empirical spatial autocorrelation without introducing topological artifacts near the medial wall. Detailed spatial similarity metrics are provided in the **Supplementary Materials**.

### Effective connectivity analysis

*Regression DCM.* To investigate directed signal flow, we applied rDCM implemented in TAPAS (version 6.0.1; https://www.tnu.ethz.ch/de/software/tapas; (Frässle, Aponte, et al., 2021)). In a nutshell, conventional DCM infers hidden dynamics by modeling the brain as a deterministic network where neural states are coupled via directed connections and translated into macroscopic BOLD signals through a biophysical hemodynamic forward model (Friston et al., 2003). rDCM reformulates DCM with the following modifications: *(i)* state and observation equations are expressed in the frequency domain (Frässle et al., 2017), *(ii)* the nonlinear hemodynamic model is replaced by a linear hemodynamic response function, *(iii)* a mean-field approximation assumes independence of parameters across regions, and *(iv)* conjugate priors are imposed on noise precision and connectivity. Through these adaptations, rDCM operates as an efficient Bayesian linear regression, enabling the estimation of directed connectivity across large-scale networks.

We utilized a spatially targeted approach guided by our primary FC findings. The cerebral cortex was parcellated using the Schaefer 400-region atlas (Schaefer et al., 2018) mapped to fsLR 10k. For each subcortical subdivision (8 thalamic and 10 striatal), we extracted the directed connectivity parameters, comprising Subcortex-to-Cortex (S→C) and Cortex-to-Subcortex (C→S) efferent and afferent pathways, restricted to the cortical parcels that fell within the previously established positive and negative FC significance masks. This approach suggests whether the FC hyper- or hypo-connectivity was driven by ascending (S→C) or descending (C→S) flow.

### Data harmonization

ComBat harmonization (Fortin et al., 2017) was applied to preserve biological variance associated with diagnostic groups (ASD, ADHD, TD), age, and sex while controlling for site effects.

### Statistical between-group comparisons

Statistical modeling was conducted using surface-based linear models in BrainStat (http://brainstat.readthedocs.io) (Larivière et al., 2023).

#### Surface-Based Comparisons

Our primary analysis, as shown in **Figure 1**, focused on direct comparison between clinical cohorts (ASD *vs*. ADHD). To quantify the magnitude of connectivity alterations, we computed vertex-wise Cohen’s d effect size maps for all group contrasts. Multiple comparison correction was performed via RFT with a cluster-defining threshold of p < 0.001 and a FWE rate < 0.05. These results identified robust positive and negative significance masks that constrained all subsequent analyses.

**Figure 1.**
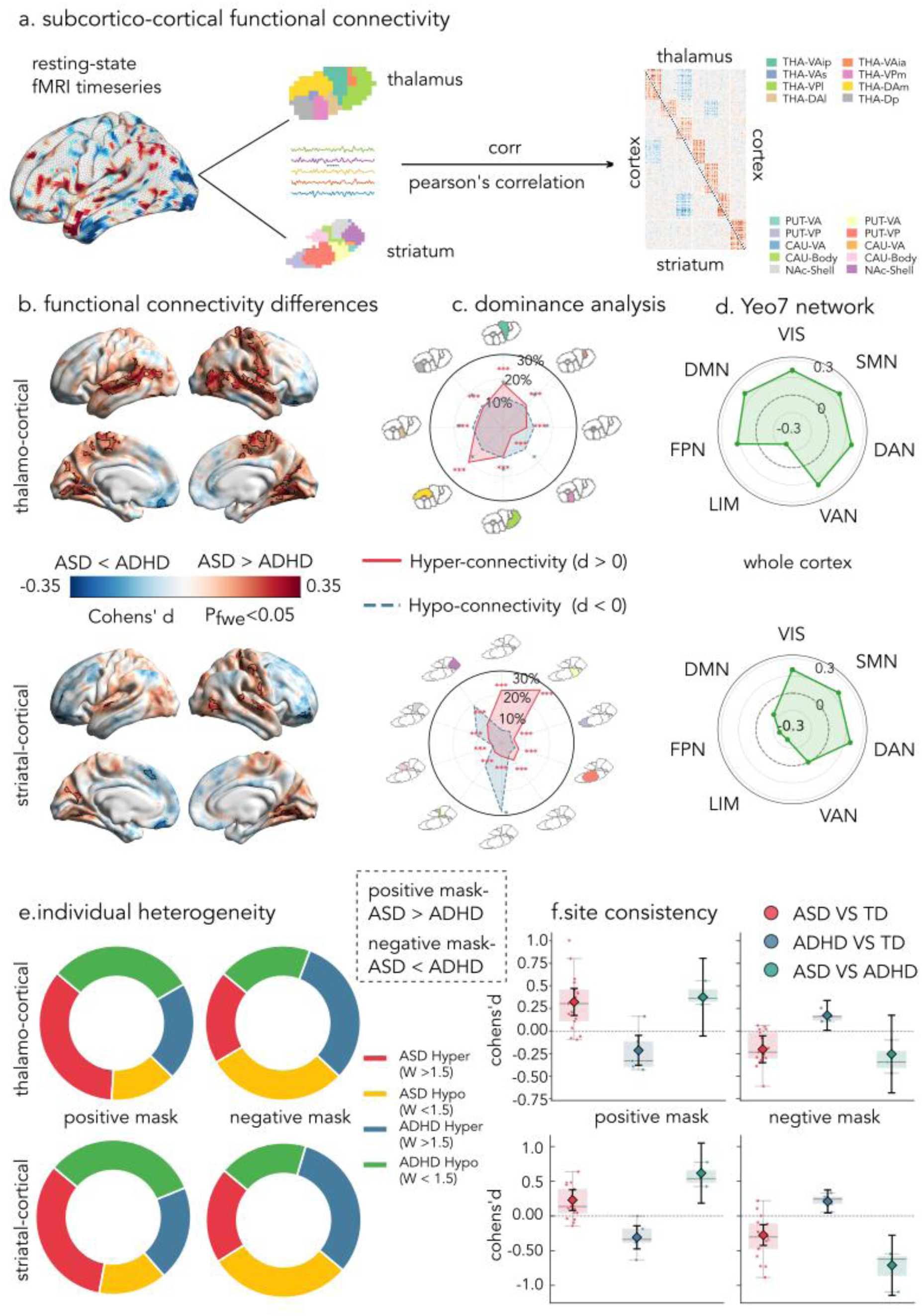
Diverging functional connectivity alterations of subcortical-cortical circuits in ASD and ADHD. a. Schematic representation of the surface-based analytical pipeline for subcortical-cortical FC. **b.** Vertex-wise Cohen’s d effect size maps contrasting the ASD and ADHD groups (cluster P_FWE_ < 0.05, cluster-defining threshold p < 0.001). The positive/red mask denotes greater connectivity in ASD (ASD > ADHD), while the negative/blue mask indicates greater connectivity in ADHD (ADHD > ASD). Solid black lines delineate the boundaries of significant clusters. **c.** Subdivisional dominance analysis illustrating the proportional contribution (%) of individual thalamic and striatal subdivisions to the significant cortical alterations. Asterisks denote statistical significance derived from 5,000 permutations (* FDR-corrected p < 0.05, ** < 0.01, *** < 0.001). Full anatomical definitions for all thalamic and striatal subdivisions derived from the Tian atlas (*e.g.,* VAip, Pvpr) are detailed in Supplementary Table S4. **d.** Functional topography mapped onto the Yeo 7-network parcellation. The radar plots display the mean Cohen’s d values extracted from the significant whole cortex. The bold inner dashed line represents zero effect size. **e.** Proportional breakdown of individualized pathological deviations (|W| > 1.5) underlying the group-level FC differences, derived from normative deviation mapping against the TD baseline. **f.** Cross-site consistency of the observed FC alterations. Box-and-swarm plots show Cohen’s d values for each independent scanning site (n ≥ 5). The solid diamonds with error bars represent the pooled meta-analytical effect sizes and their 95% confidence intervals, respectively.

### Statistical Modeling of Effective Connectivity

Consistent with the FC analysis, models were adjusted for age and sex, with diagnostic groups as the primary contrast of interest (*e.g*., ASD vs. ADHD). Statistical significance for the S→C and C→S alterations was determined using the FDR procedure, thresholded at p < 0.05 (Benjamini & Hochberg, 1995).

### Individualized deviation mapping

To capture individual-level heterogeneity, we mapped individualized deviations of subcortical-cortical FC relative to the large TD baseline cohort using standardized W-scores, based on established normative deviation frameworks (Marquand et al., 2016; Xie et al., 2024). Age and sex were regressed out to establish a typical neurodevelopmental trajectory. For every participant, deviations from this normative baseline were quantified using W-scores. We calculated the frequency of positive (W > 1.5) and negative (W < -1.5) deviations within the corresponding positive and negative significance masks identified in the primary group comparisons. To next evaluate individual-level heterogeneity in directed signal flow, we extended this W-score framework to the rDCM parameters. Using the TD cohort as a baseline, individualized W-scores were computed for both S→C and C→S connections. The frequencies of extreme bidirectional deviations (|W| > 1.5) were then quantified to elucidate the predominant directionality of these deviations.

### Multivariate brain-behavior associations

To enable unified transdiagnostic evaluation, disparate clinical instruments (*e.g.,* ADOS, SRS) were Z-normalized within their respective cohorts. The harmonized phenotypic battery (Y-matrix) encompassed generalized cognitive indices (Full-Scale, Verbal, and Performance IQ) and domain-specific symptom severities (inattention, hyperactivity, social deficits, and restricted/repetitive behaviors).

The neuroimaging X-matrix concatenated individualized W-scores for vertex-wise FC and parcel-wise rDCM parameters, extracted strictly from the diagnosis-specific significance masks (ASD-*vs*-TD and ADHD-*vs*-TD). We applied an asymmetric confound regression strategy. The behavioral scores Y-matrix was residualized against age, age², sex, mean FD, and scanning site. In contrast, because W-scores intrinsically account for normative demographic trajectories and site effects (via prior ComBat harmonization), the X-matrix was residualized against mean FD to prevent statistical over-correction.

To map high-dimensional brain features to clinical profiles, we employed a Principal Component-Partial Least Squares (PC-PLS) framework (Krishnan et al., 2011) across the transdiagnostic cohort and within diagnosis-specific subsets. To reduce the number of parameters, PCA was first applied to the residualized X-matrix. The retained brain components and residualized Y-matrix were entered into a PLS regression to identify orthogonal latent variables (LVs) maximizing brain-behavior covariance. The significance of the extracted LVs was evaluated via 5,000 subject-level permutations, wherein the rows of the residualized behavioral matrix were randomly shuffled to establish a null distribution of brain-behavior covariance. Finally, 5,000 bootstrap resampling iterations were performed to calculate Bootstrap Ratios (BSR), identifying the most stable neural features driving the neurobehavioral axes (McIntosh & Lobaugh, 2004).

## Results

### Differential FC alterations in subcortical-cortical circuits in ASD and ADHD

To delineate the functional architecture of subcortical-cortical circuits in ASD and ADHD **(Figure 1a)**, we computed vertex-wise linear models that compared FC patterns between our study cohorts (P*_FWE_* < 0.05; **Figure 1b)**. Specifically, thalamocortical connectivity was elevated in ASD relative to ADHD (Positive mask: peak Cohen’s d = 0.330) across widespread sensorimotor and associative cortices, with ADHD showing only sparse localized hyperconnectivity in the medial orbitofrontal cortex (Negative mask: peak d = -0.309). Conversely, striatocortical comparisons revealed higher connectivity in ASD (peak d = 0.309) across posterior sensory regions, while ADHD exhibited stronger striatal coupling primarily in prefrontal areas (peak d = -0.317). We also compared both diagnostic groups against typically developing (TD) controls **(Supplementary Figure S1)**.

Dominance analysis **(Figure 1c)** quantified contributions of thalamic and striatal subdivisions. ASD-related thalamic increases were broadly supported by all eight subdivisions (P*_FDR_* < 0.001), primarily led by the dorsoanterior (Dam, 20.1%) and ventroanterior (VAip, 18.0%) complexes. In the striatocortical network, the heightened connectivity in ADHD was singularly accounted for by the caudal dorsoanterior striatum (Cdar, 28.4%; P*_FDR_* < 0.05). In contrast, the elevated striatal coupling in ASD was diffusely supported by all ten subdivisions, largely dominated by the posterior putamen (Pdar, 27.3%; Pvar, 21.9%).

Mapping these alterations onto the Yeo-Krienen seven network assignment **(Figure 1d)** highlighted distinct topographical profiles. ASD-related increases were broadly distributed across sensory-associative systems, predominantly mapping to the visual (peak d = 0.214), default mode, and somatomotor networks. In contrast, ADHD-related effects were preferentially localized to higher-order limbic (peak d = -0.218) and executive networks.

Normative deviation mapping against a TD baseline quantified individual-level pathological deviations (|W| > 1.5; **Figure 1e**). In both thalamic and striatal systems, regions exhibiting ASD > ADHD connectivity at the group level were primarily driven by a high relative burden of hyperconnectivity in ASD (33.2%–35.1% of total deviations) and hypoconnectivity in ADHD (30.5%–32.7%; chance = 25%). This divergence was similarly robust in regions showing the reverse pattern (ADHD > ASD), where ADHD individuals exhibited pronounced hyperconnectivity contributions (31.4%–31.5%) alongside marked functional reductions in ASD (29.5%–29.9%).

Analysis across all imaging sites confirmed the robustness of these effects **(Figure 1f).** Crucially, the differences between ASD and ADHD were driven by opposite deviations from typical development. For example, within the striatocortical network, individuals with ADHD exhibited elevated connectivity relative to the normative baseline (d = 0.210), whereas individuals with ASD showed reduced connectivity (d = -0.275). This divergence created a large contrast between the two clinical groups (pooled ADHD > ASD d = -0.710, 95% CI [-1.144, -0.277]). This pattern of opposing deviations—where one condition shows hyperconnectivity and the other hypoconnectivity—was consistently observed across all major subcortical-cortical networks.

### Diverging signal flow patterns in ASD and ADHD

To infer directed signal flow underpinning observed FC alterations, regression dynamic causal modeling (rDCM) estimated Subcortex-to-Cortex (S→C) and Cortex-to-Subcortex (C→S) effective connectivity **(Figure 2a)**. Group comparisons within the FC-derived significance masks (FDR< 0.05; Figure 2b) revealed distinct directional alterations. In the ASD-hyperconnected thalamocortical network, elevated FC was supported by enhanced bidirectional connectivity in ASD, distributed evenly across S→C and C→S paths **(Figure 2b, top left)**. Peak enhancements originated from the anterior intralaminar nucleus (S → C peak d = 0.289) and targeted the anteromedial thalamus (C→S peak Cohen’s d = 0.268). Within ADHD-hyperconnected thalamic regions, the ASD group exhibited selectively reduced S→C flow (peak Cohen’s d = -0.158; **Figure 2b, top right**). In the ASD-hyperconnected striatocortical network, elevated effective connectivity in ASD involved 26 bilateral pathways **(Figure 2b, bottom left)**. This was characterized by enhanced C→S flow targeting the accumbens core (peak d = 0.260) and anterior putamen, alongside elevated S→C flow from the dorsal caudate. In contrast, the ADHD-hyperconnected striatal network was mirrored by heightened flow in the ADHD cohort, primarily driven by enhanced C→S flow to the caudate body (peak Cohen’s d = -0.232) and increased S→C flow from the caudate tail **(Figure 2b, bottom right).**

**Figure 2.**
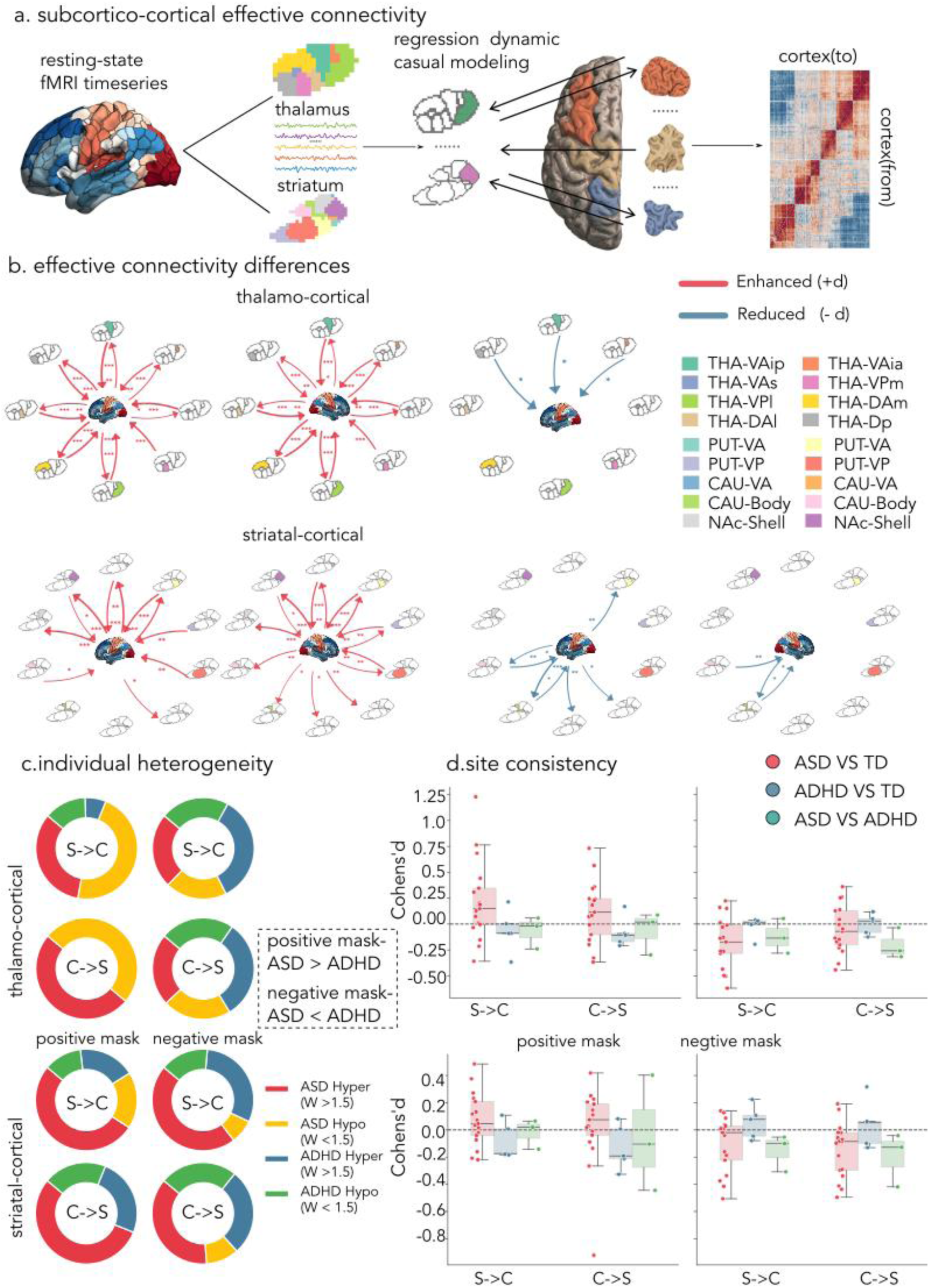
Directional architecture and individual-level drivers of subcortical-cortical dysregulation. **a.** Schematic representation of the rDCM analytical framework. **b.** Group-level effective connectivity alterations. Hub- and-spoke network plots display significant Subcortex-to-Cortex (S→C) and Cortex-to-Subcortex (C→S) pathways within the established Positive and Negative FC masks (see Figure 1). Line thickness scales with the magnitude of the effect size (Cohen’s *d*). Red lines denote enhanced modulation, whereas blue lines indicate reduced modulation relative to the comparison group. Asterisks denote statistical significance derived from surface-based linear models (* FDR-corrected p < 0.05, ** < 0.01, *** < 0.001). **c.** Proportional breakdown of extreme individualized directional deviations. The 2x2 donut charts quantify the frequency of extreme hyper- or hypo-signaling events (|W| > 1.5) for both S→C and C→S pathways, derived from normative deviation mapping against the TD baseline. **d.** Cross-site consistency of macroscopic causal flows. Box-and-swarm plots illustrate the aggregated Cohen’s *d* values for overall S→C and C→S macro-circuits across independent scanning sites, contrasting the clinical groups against the TD baseline.

Normative deviation mapping evaluated the proportional contribution of extreme effective connectivity states (|W| > 1.5) **(Figure 2c)**. In the ASD-hyperconnected thalamocortical network, extreme deviations were overwhelmingly driven by the ASD group, accounting for 80.0% of thalamo-cortical (S→C) and 100.0% of cortico-thalamic (C→S) extreme deviations. Notably, these deviations reflected a heterogeneous mix of hyper-and hypo-connectivity (S→C: 33.3% hyper vs. 46.7% hypo; C→S: 50.0% hyper vs. 50.0% hypo). Within striatocortical networks, extreme deviations were predominantly characterized by hyper-connectivity. Specifically, in ASD-biased striatal regions, these deviations were primarily driven by the ASD group showing hyper-connectivity in both the striato-cortical (S→C; 52.0%) and cortico-striatal (C→S; 55.0%) directions. In the ADHD-biased striatal network, hyper-connectivity remained the most frequent extreme deviation, with comparable contributions from both the ASD and ADHD groups. To contextualize these directional disruptions against typical neurodevelopment, subdivision-level parameters were aggregated into global macro-circuit averages (overall S→C and C→S flows) for multi-site analysis (**Figure 2d**). In the ASD-hyperconnected thalamocortical network, macroscopic S→C flow was significantly elevated in ASD relative to TD (pooled Cohen’s d = 0.221, 95% CI [0.019, 0.423]), whereas ADHD remained statistically indistinguishable from normative levels. Conversely, in ADHD-hyperconnected thalamic regions, ASD demonstrated significantly reduced macroscopic S→C flow relative to TD (pooled Cohen’s d = -0.180). In the striatocortical system, global deviations from the baseline were concentrated within the ADHD-biased network. This macroscopic divergence was primarily attributable to the ASD group, which exhibited widespread global reductions in both S→C driving (pooled Cohen’s d = -0.107) and C→S feedback (pooled Cohen’s d = -0.149) flows relative to healthy controls, while the ADHD group maintained normative macro-circuit flows. Within the ASD-biased striatal network, directional macro-flows were largely normative. Collectively, these directed analyses indicate that ASD-related alterations were characterized by widespread bidirectional thalamocortical dysregulation alongside reduced striatocortical macro-flows, whereas ADHD-related alterations were more selectively localized to focal striatal regulatory circuits.

### Multivariate brain-behavior associations and clinical specificity

PLS analysis integrated FC and rDCM features with behavioral phenotypes across ASD and ADHD **(Figure 3a)**. In the transdiagnostic cohort **(Figure 3b)**, the PLS model identified a significant latent variable (Permutation *p* = 0.015, r = 0.110) characterized by a negative association with cognitive performance, primarily weighted by Full Scale IQ (Loading = -0.987). The associated neural profile comprised 62 stable features (Bootstrap Ratio |BSR| ≥ 1.96). Neural correlates were localized entirely within the striatocortical circuitry (100%), with no significant contributions from thalamic networks. Furthermore, this pattern mainly loaded on effective flow, representing 75.8% of the selected features (40.3% C→S; 35.5% S→C). All 62 neural features emerged from the ASD-hyperconnected striatal networks.

**Figure 3.**
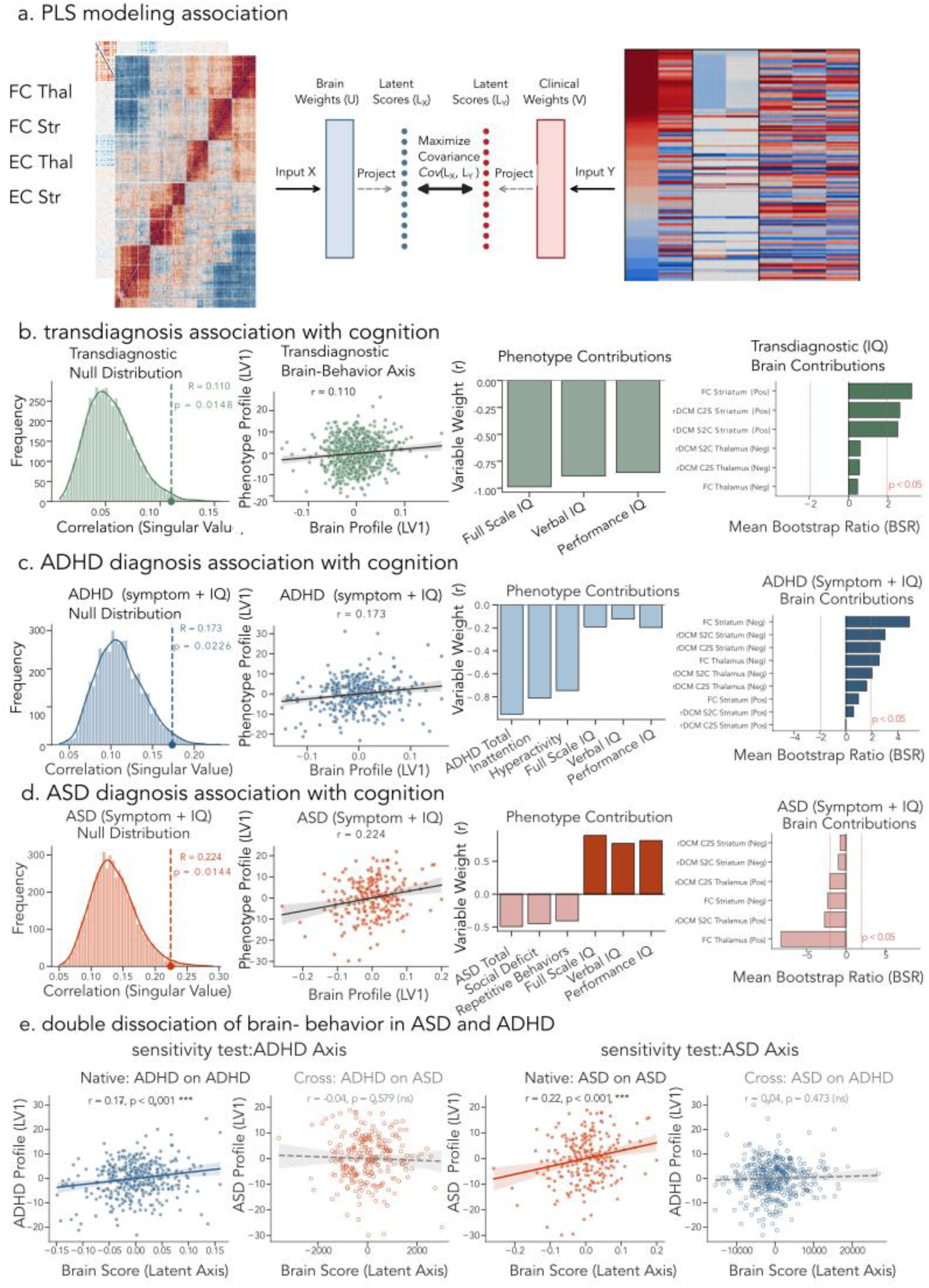
Multivariate brain-behavior associations and transdiagnostic clinical specificity. **a.** Schematic illustration of the PLS analytical framework, integrating high-dimensional functional (FC) and effective (rDCM) connectivity features with clinical and cognitive assessments. **b.** Transdiagnostic PLS analysis within the combined cohort. Sub-panels display the permutation null distribution of singular values (5,000 iterations), the scatter plot of individual brain and cognitive latent variable (LV) scores, the variable weights (r) for behavioral contributions, and the Bootstrap Ratios (BSR) for the top neural drivers. Red dashed lines denote the statistical significance threshold (|BSR| > 1.96). **c.** Diagnosis-specific PLS analysis in the ADHD cohort, detailing the permutation distribution, LV score correlations, phenotype contributions (weighted toward core ADHD symptom severity), and corresponding top sub-nuclei neural drivers. **d.** Diagnosis-specific PLS analysis in the ASD cohort, presenting the corresponding permutation distribution, LV scores, phenotype contributions (weighted predominantly toward cognitive performance), and top neural drivers. **e.** Transdiagnostic projection and double dissociation. The 2x2 scatter plots contrast the predictive performance of the native brain axes within their own cohorts (left column) against their out- of-sample projections onto the alternate cohort (right column). The statistical significance of the dissociations between native and projected correlation coefficients was evaluated using Fisher’s Z-transformation.

PLS analysis within ADHD **(Figure 3c)** resulted in one significant latent axis (Permutation *p* = 0.023, r = 0.173) weighted towards clinical symptoms, led by overall severity (Loading = -0.950) and inattention (Loading = -0.812), with minimal contribution from cognitive measures. The neural profile comprised 265 stable features (|BSR| ≥ 1.96), balanced between the thalamocortical and striatocortical networks. All selected features exhibited positive weights and were anchored within the ADHD-hyperconnected functional networks. These multivariate associations were supported by a mixture of static FC and balanced bidirectional effective signaling, with the strongest contributions emerging from striatal static FC and S→C flow. In the ASD cohort **(Figure 3d)**, the PLS model identified a latent axis (Permutation *p* = 0.014, r = 0.224) showing a profile distinct from the ADHD model. Rather than core symptomatology, the behavioral axis was primarily weighted towards IQ measures (Full Scale IQ Loading = 0.894). The associated neural profile comprised 2,198 stable features. In contrast to the ADHD model, these cognitively weighted neural features were characterized almost entirely by negative weights (>99% with BSR < 0) and were localized to the thalamocortical network (>97%). This pattern was concentrated within the ASD-hyperconnected thalamic network, dominated by static thalamocortical FC and altered S→C flow from the thalamus.

To test the clinical specificity of these diagnosis-specific neural axes, a transdiagnostic projection analysis was conducted (**Figure 3e**). Within the ADHD discovery cohort, the primary PLS latent variable captured an in-sample brain–behavior covariation pattern (r = 0.173, p < 0.001). To evaluate whether this neural axis was specific to ADHD or transdiagnostically generalizable, we fixed the spatial weight matrix of this ADHD neural axis and applied it to the connectivity data of the independent ASD cohort, thereby generating out-of-sample brain scores for ASD individuals. Crucially, these projected brain scores failed to track the corresponding behavioral scores within the ASD cohort (r = -0.037, p = 0.579). A Fisher Z-transformation confirmed a significant difference between these two correlations (Z = 2.54, p = 0.011), establishing the clinical specificity of the ADHD neural profile.

Similarly, the discovery PLS model within the ASD cohort identified an in-sample latent dimension linking brain features to cognitive scores (r = 0.224, p < 0.001). When the spatial weights defining this cognitively anchored ASD brain axis were locked and projected onto the independent ADHD cohort, the correlation with cognitive performance dissolved (r = 0.037, p = 0.473). This diagnostic dissociation was statistically significant (Fisher Z = 2.29, p = 0.022). Site-level evaluations confirmed that the within-diagnosis latent correlations were consistent across scanning sites (**Supplementary Figure S3**). Together, these analyses demonstrated that ADHD-related network alterations were preferentially associated with symptom severity, whereas ASD-related connectivity profiles reflected cognitive performance.

## Discussion

The present study targeted the functional organization of subcortico-cortical systems to identify shared and distinct connectivity features in ASD and ADHD. First, FC analyses revealed preferential thalamocortical alterations in ASD and striatocortical disruptions in ADHD. Second, effective connectivity analyses based on rDCM demonstrated that these alterations were accompanied by distinct patterns in directional signal flow that went beyond symmetric atypical coupling. Specifically, within thalamocortical loops, ASD-related hyperconnectivity was reflected by coordinated bidirectional amplification—anchored by ascending drive from the anterior intralaminar nucleus and descending feedback to the anteromedial thalamus—alongside macro-circuit elevations in ascending (S→C) causal flow relative to typical development. In contrast, ADHD-related alterations dissociated along specific striatal pathways, characterized by selective cortico-striatal input to the caudate body and altered striato-cortical outflow from the caudate tail, alongside an unexpected macroscale suppression of striatocortical signal flow in ASD. Finally, multivariate brain-behavior analyses related these directed network alterations to dissociable cognitive and symptom profiles. Across both conditions, atypical striatocortical coupling was associated with lower general cognitive performance, supporting a shared role of striatocortical circuitry in overall cognitive processing. ADHD-related directional alterations furthermore tracked hyperactive and inattentive symptom severity, whereas ASD-related connectivity profiles were more strongly associated with overall IQ measures. The latter relationships critically failed to generalize when projected across cohorts, demonstrating condition-specific brain–behavior coupling. These findings demonstrate that beyond the phenotypic separations captured by static functional connectivity, resolving directional network dynamics unveils specific circuit-level and condition-specific neurodevelopmental vulnerabilities, supporting an important role of directed subcortico-cortical interactions in atypical brain development.

By interrogating cortical and subcortical systems at a subregional level, our framework enabled the identification of localized anatomical divergences between ASD and ADHD. FC dominance and rDCM analyses revealed that ASD-related dysregulation was broadly distributed across thalamic subdivisions, with prominent involvement of anterior and intralaminar subdivisions (*e.g.,* VAip, Dam). Our framework also localized thalamocortical signal flow changes in divisions previously implicated in arousal regulation and attentional gating (Sherman, 2016). Although the present study did not directly assess sensory processing, these findings are broadly consistent with models proposing atypical thalamocortical signaling in sensory over-responsivity in autism (Marco et al., 2011; Wagner et al., 2023), as well as recent work linking altered thalamocortical coupling to aversive sensory phenotypes (Hwang et al., 2022; Karavallil Achuthan et al., 2023). These findings align with resting-state and task-based fMRI studies demonstrating atypical functional architecture and recruitment of these specific thalamic subdivisions in ASD (Di Martino et al., 2014; Green et al., 2017; Nair et al., 2015). Prior task-based work has related disrupted sensory and attentional modulation to altered thalamocortical signaling in autistic individuals (Green et al., 2017; Hwang et al., 2022), supporting an empirical basis for the subcortical-to-cortical dysregulation observed here. Crucially, multivariate PLS models extended these results by showing that widespread thalamocortical dysregulation preferentially maps onto Full Scale IQ rather than core symptoms. Elevated thalamocortical signaling was inversely associated with generalized cognitive performance in ASD. As static resting-state fMRI data precludes direct mechanistic interpretations, it remains to be established whether the observed negative associations align with theoretical frameworks proposing that atypical thalamo-cortical interactions impact large-scale network coordination during higher-order cognition in autism (Dinstein et al., 2012; Gomot & Wicker, 2012; Halassa & Kastner, 2017; Lawson et al., 2014; Mottron et al., 2006). Such atypical signal flow may also impact communication between sensory and transmodal systems, and contribute to sensory and perceptual alterations in autism (Bernhardt et al., 2025; Hong et al., 2019; Mottron et al., 2006; Park et al., 2021).

In contrast, the neurobiological architecture of ADHD was prominently weighted toward striatocortical alterations. Unlike the more diffuse pattern observed in ASD, ADHD-related striatal deviations were primarily localized to the caudate nucleus, a region previously implicated in cortico-striatal circuits supporting action selection and inhibitory control (Graybiel, 2008). Within these networks, our rDCM analyses revealed widespread disruptions in directed signaling, involving both altered cortico-striatal (C→S) and elevated striato-cortical (S→C) effective connectivity. This pattern is consistent with recent evidence suggesting that enhanced directed cortico-striatal signaling may relate to hyperactive and impulsive symptom dimensions (Zhang et al., 2024). Indeed, multivariate modeling indicated that these striatocortical disruptions were preferentially associated with core ADHD symptoms (inattention and hyperactivity), rather than generalized cognitive performance. While the thalamus is increasingly recognized as an integrative hub for cognitive control (Hwang et al., 2017; Hwang et al., 2022; Shine et al., 2019; Shine et al., 2023), our results further suggest a prominent contribution of striatal pathways to motor and attentional regulation in ADHD. Furthermore, positive weights in the PLS model are consistent with a pattern of increased subcortical–cortical coupling, a network configuration that has been hypothesized to relate to behavioral disinhibition (Hong, 2023; Mills et al., 2012). By mapping directed circuit alterations onto symptom severity, these findings suggest that atypical signal flow across striatal–cortical circuits tracks clinical ADHD symptoms, while showing a weaker relationship with general intellectual performance.

The application of an individualized analysis framework to both functional and effective connectivity parameters provided additional insight into patient heterogeneity. Traditional case-control analyses inherently rely on group means, which can obscure the elevated inter-individual heterogeneity seen across neurodevelopmental conditions (Feczko et al., 2019; Lombardo et al., 2019; Wolfers et al., 2020). By estimating individualized W-scores relative to a typically developing reference distribution(Xie et al., 2024), we show that macroscopic network alterations in ASD and ADHD do not reflect uniform group-level shifts, but rather distributed patterns of deviation across a shared subcortical–cortical system. Within the thalamocortical network, a subset of individuals with ASD exhibited pronounced hyperconnectivity, whereas individuals with ADHD showed a tendency toward reduced connectivity relative to the normative baseline. Importantly, even in cases where a similar direction of deviations was observed, such as elevated striatal coupling across both cohorts, our multivariate models indicate that these alterations are associated with distinct clinical profiles. In ASD, these associations more closely point to atypical overall cognitive performance, while they point more strongly to symptom expression in ADHD. While our region-of-interest definitions were guided by group-level statistical masks, these findings underscore the value of subsequent individualized connectomic profiling for capturing the full spectrum of neurobiological variation in neurodevelopmental indications. In this context, normative gradient modeling frameworks pioneered by Marquand and colleagues (Haak et al., 2018; Marquand et al., 2016; Rutherford et al., 2022) provide an ideal avenue to chart how individual deviations express along continuous topographical axes.

Several methodological considerations warrant discussion. First, aggregating multi-site cohorts (ABIDE, ADHD-200, and HBN) introduces technical heterogeneity, including variations in scanner platforms, field strengths, and acquisition protocols. To mitigate these batch effects, we applied ComBat harmonization while preserving biological variance of interest (age, sex, and diagnosis). Crucially, site-stratified analyses confirmed consistent effect directions across scanning sites, indicating that macro-circuit alterations capture robust biological patterns rather than acquisition artifacts. Second, while rDCM provides an efficient Bayesian framework for estimating effective connectivity from BOLD time series, it remains only a proxy of underlying neural dynamics (Frässle, Harrison, et al., 2021; Frässle et al., 2017; Stephan et al., 2010). Future studies leveraging higher-temporal-resolution modalities, such as MEG or EEG, will be important to further validate the inferred directionality of signal flow, although these electrophysiological techniques remain challenged in mapping signals from deep subcortical and mesiotemporal structures (Attal & Schwartz, 2013; Baillet, 2017; Pizzo et al., 2019). Thirdly, integrating functional MRI analysis approaches with multimodal information, in particular microstructural profiling of cortical and mesiotemporal architectures (Paquola et al., 2020; Paquola et al., 2025), may offer a promising avenue to contextualize and constrain these macroscale directed interactions. Specifically, linking subcortico-cortical dynamics to distinct cytoarchitectonic and receptor profiles such as the balance between thalamic “core” and “matrix” populations and their alignment along sensory-to-transmodal cortical hierarchies (Müller et al., 2020; Shine et al., 2023), could provide valuable cellular-level grounding for the directional circuit alterations identified here. Finally, the cross-sectional nature of the present cohort limits direct inferences regarding individual-level developmental change over time. Future longitudinal connectomic studies (Mills et al., 2016) and large-scale normative deviation frameworks may address subcortical–cortical connectivity changes robustly across the lifespan (Bethlehem et al., 2022; Rutherford et al., 2022), and may chart how these circuit-level vulnerabilities emerge and diverge across neurodevelopment. In conclusion, this study delineates the functional, directional, and clinical organization of subcortical–cortical circuits in ASD and ADHD. By integrating large-scale neuroimaging with normative deviation mapping and multivariate behavioral mapping, we identify both a shared striatocortical vulnerability and a robust pattern of clinical divergence between the two conditions. Specifically, the widespread thalamocortical dysregulation and altered ascending signaling observed in ASD were primarily associated with overall indices cognitive performance, rather than core autistic symptoms. Conversely, ADHD was characterized by more pronounced subcortical involvement, with stronger contributions from striatal regulatory circuits that were preferentially associated with inattention and hyperactivity. Together, these findings provide a transdiagnostic framework linking directional circuit-level alterations to distinct patterns of clinical and cognitive variation across neurodevelopmental conditions.

## Acknowledgments

G.Z. is funded by the China Scholarship Council. A.C.E acknowledges funding from CFREF/HBHL (252428) and CIHR PJT-173236 (254573). B.C.B. acknowledges research support from the National Science and Engineering Research Council of Canada (NSERC RGPIN-2025-05932), CIHR (FDN-154298, PJT-174995, PJT-191853, PJT-203761), SickKids Foundation (NI17-039), Helmholtz International BigBrain Analytics and Learning Laboratory (HIBALL), HBHL, Brain Canada Foundation, FRQS, Tier-2 Canada Research Chairs Program, and The Centre for Excellence in Epilepsy at the Neuro (CEEN).

## Supplementary Information

**Supplementary Figure S1.**
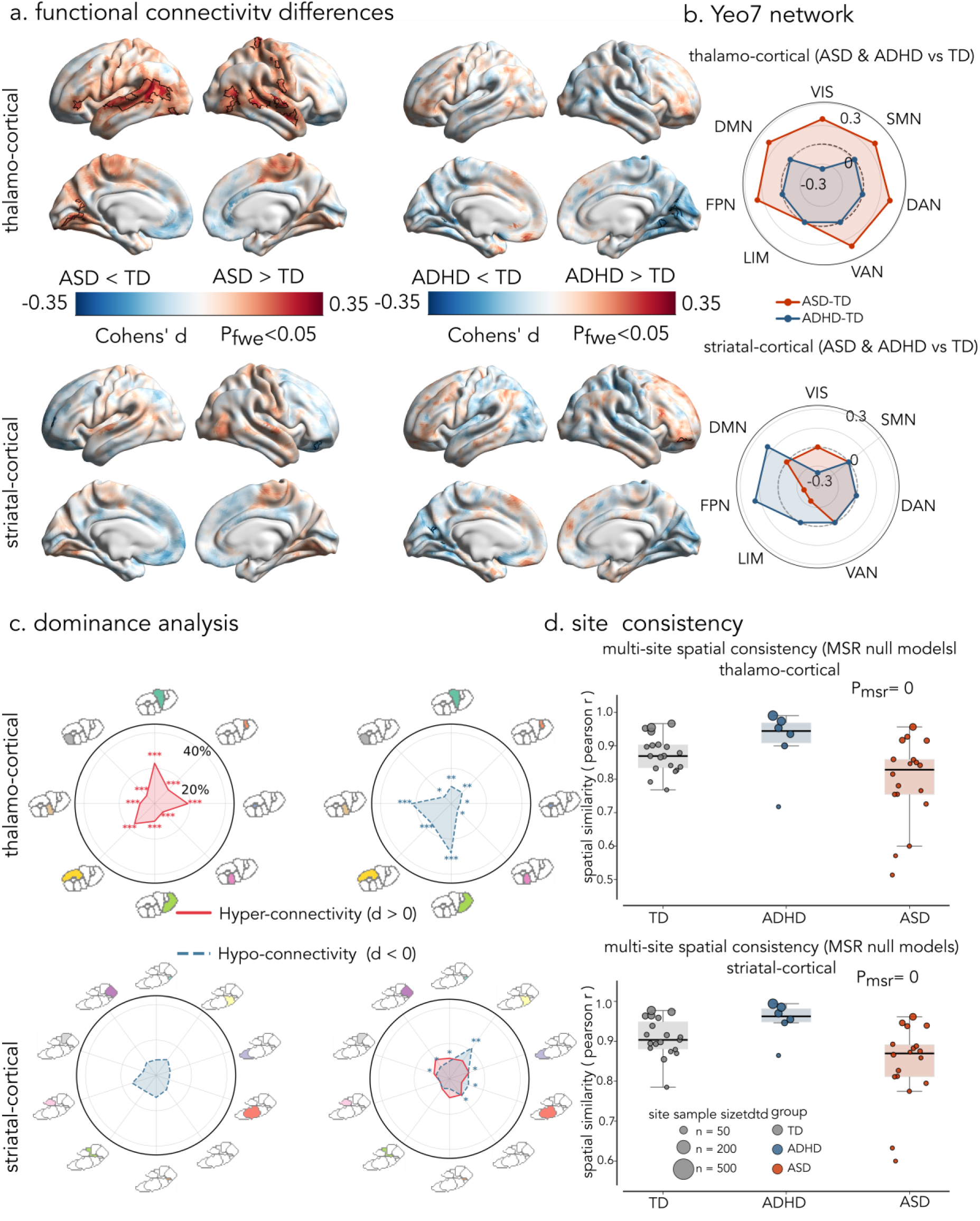
Baseline-contrasted functional architecture of subcortical-cortical circuits in ASD and ADHD. **a.** Vertex-wise Cohen’s d effect size maps contrasting the ASD and ADHD groups separately against the typically developing (TD) baseline (cluster P_FWE_ < 0.05, cluster-defining threshold p < 0.001). positive/red denotes hyper-connectivity (Clinical > TD), while negative/blue denotes hypo-connectivity (Clinical < TD). Solid black lines delineate the boundaries of significant clusters. **b.** Functional topography mapped onto the Yeo 7- network parcellation. The radar plots display the whole-brain mean Cohen’s d values for the ASD vs. TD and ADHD vs. TD contrasts. The bold inner dashed line represents zero effect size. **c.** Subdivisional dominance analysis illustrating the proportional contribution (%) of individual thalamic and striatal subdivisions to the significant cortical alterations defined by the respective disorder-specific FC masks in (a). Asterisks denote statistical significance derived from 5,000 permutations (* FDR-corrected p < 0.05, ** < 0.01, *** < 0.001). Full anatomical definitions for all subcortical subdivisions (e.g., VAip, Pvpr) are detailed in Supplementary Table S4. **d.** Cross-site consistency and spatial robustness of the FC profiles across the ASD, ADHD, and TD cohorts. Statistical significance of the spatial consistency was evaluated against spatial null models generated via Moran Spectral Randomization (MSR) to rigorously account for spatial autocorrelation.

**Supplementary Figure S2.**
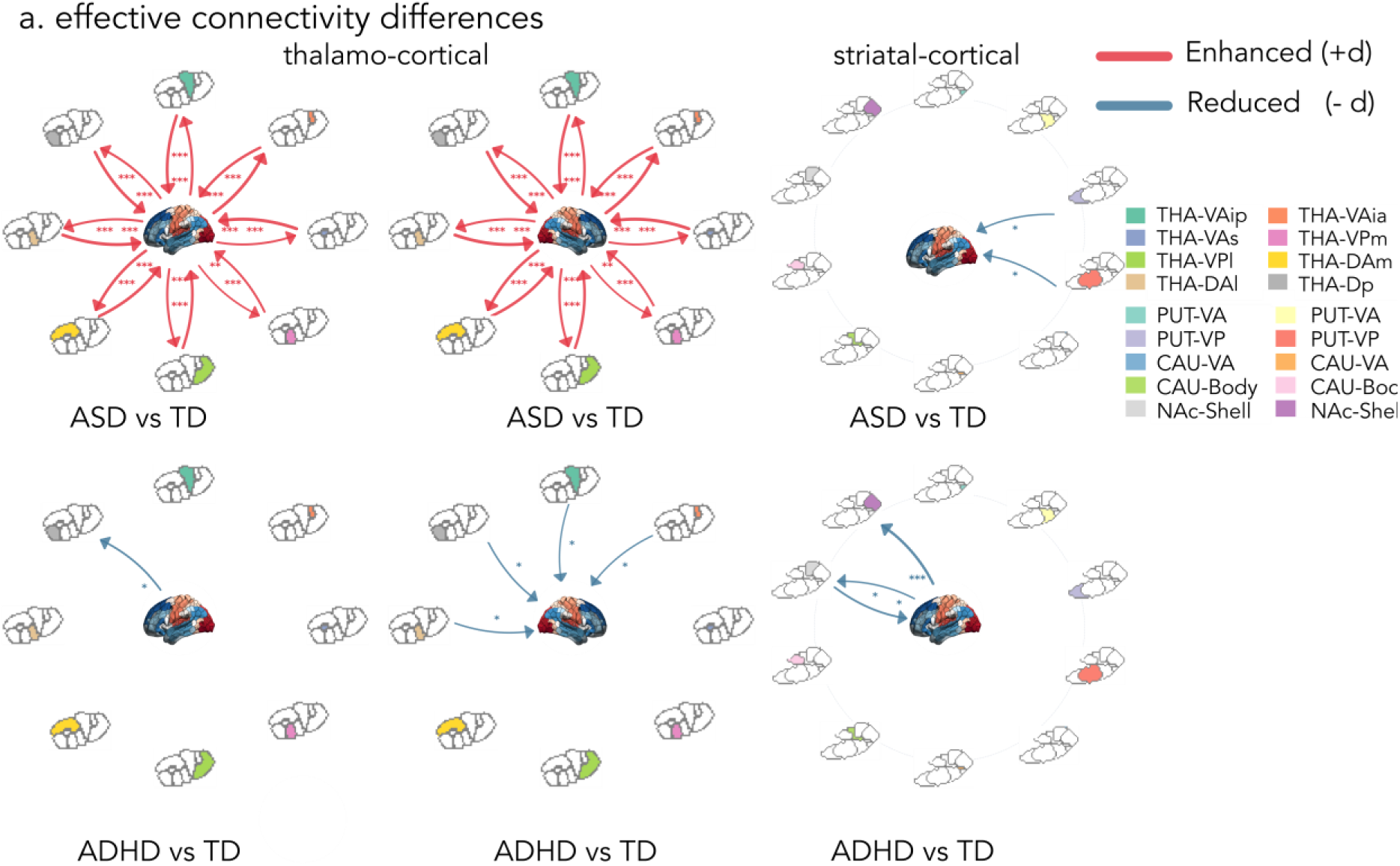
Baseline-contrasted effective connectivity alterations of subcortical-cortical circuits in ASD and ADHD. Hub-and-spoke network plots display the significant Subcortex-to-Cortex (S→C) and Cortex-to-Subcortex (C→S) pathways for the ASD and ADHD cohorts when contrasted separately against the typically developing (TD) baseline. Line thickness scales with the magnitude of the effect size (Cohen’s d). Red lines denote enhanced modulation, whereas blue lines indicate reduced modulation relative to the TD group. Asterisks denote statistical significance derived from surface-based linear models (* FDR-corrected p < 0.05, ** < 0.01, *** < 0.001).

**Supplementary Figure S3.**
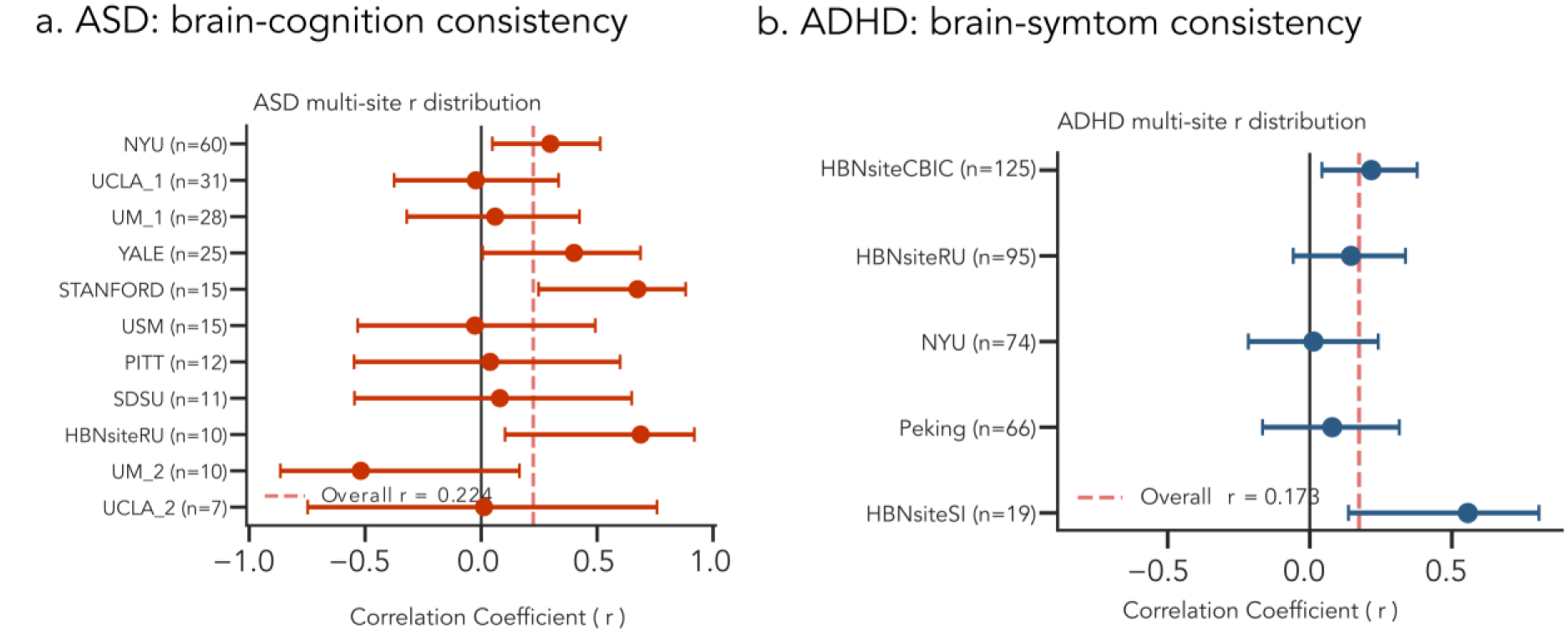
Cross-site robustness of the multivariate brain-behavior associations in ADHD and ASD. Forest plots illustrating the consistency of the within-diagnosis latent correlations derived from the Partial Least Squares (PLS) analyses across independent scanning sites. (a) Site-specific correlation coefficients (r) between the ASD-derived, thalamus-dominated brain axis and generalized cognitive performance (i.e., Full-Scale IQ) within the ASD cohort. (b) Site-specific Pearson correlation coefficients (r) between the ADHD-derived composite brain axis and core clinical symptom severity (i.e., inattention and hyperactivity scores) within the ADHD cohort. For both panels, markers represent the correlation effect size for each individual site, with error bars denoting the 95% confidence intervals. The overall pooled correlation for each cohort is indicated for reference, demonstrating that the multivariate clinical mappings are highly consistent and not driven by site-specific confounding effects.

**Supplementary Table S1.** Site-specific demographic and head motion profiles.

| Dataset | Site | Diagnosis | N | Age (years, Mean ± SD) | Sex (Male n, %) | Mean FD (mm, Mean ± SD) |
| --- | --- | --- | --- | --- | --- | --- |
| ABIDE | KKI | ASD | 16 | 9.92 ± 1.47 | 13 (81.2%) | 0.142 ± 0.077 |
| ABIDE | KKI | TD | 32 | 10.13 ± 1.27 | 23 (71.9%) | 0.098 ± 0.045 |
| ABIDE | LEUVEN_2 | ASD | 15 | 13.92 ± 1.31 | 12 (80.0%) | 0.117 ± 0.066 |
| ABIDE | LEUVEN_2 | TD | 20 | 14.34 ± 1.51 | 15 (75.0%) | 0.088 ± 0.027 |
| ABIDE | MAX_MUN | ASD | 6 | 11.83 ± 3.31 | 6 (100.0%) | 0.169 ± 0.050 |
| ABIDE | MAX_MUN | TD | 3 | 9.33 ± 2.08 | 3 (100.0%) | 0.080 ± 0.035 |
| ABIDE | NYU | ASD | 60 | 11.34 ± 2.69 | 54 (90.0%) | 0.091 ± 0.051 |
| ABIDE | NYU | TD | 73 | 12.45 ± 2.89 | 54 (74.0%) | 0.061 ± 0.043 |
| ABIDE | OLIN | ASD | 12 | 15.17 ± 2.48 | 11 (91.7%) | 0.136 ± 0.064 |
| ABIDE | OLIN | TD | 8 | 14.75 ± 2.25 | 7 (87.5%) | 0.136 ± 0.078 |
| ABIDE | PITT | ASD | 17 | $13.97 \pm 2.37$ | 13 (76.5%) | $0.126 \pm 0.067$ |
| ABIDE | PITT | TD | 14 | $14.37 \pm 1.74$ | 11 (78.6%) | $0.146 \pm 0.051$ |
| ABIDE | SDSU | ASD | 12 | $15.05 \pm 1.67$ | 12 (100.0%) | $0.062 \pm 0.052$ |
| ABIDE | SDSU | TD | 22 | $14.22 \pm 1.90$ | 16 (72.7%) | $0.070 \pm 0.045$ |
| ABIDE | STANFORD | ASD | 19 | $10.01 \pm 1.62$ | 15 (78.9%) | $0.101 \pm 0.062$ |
| ABIDE | STANFORD | TD | 19 | $9.89 \pm 1.62$ | 15 (78.9%) | $0.092 \pm 0.052$ |
| ABIDE | TRINITY | ASD | 13 | $14.94 \pm 1.58$ | 13 (100.0%) | $0.097 \pm 0.040$ |
| ABIDE | TRINITY | TD | 14 | $14.62 \pm 1.68$ | 14 (100.0%) | $0.085 \pm 0.052$ |
| ABIDE | UCLA_1 | ASD | 31 | $13.58 \pm 2.59$ | 26 (83.9%) | $0.097 \pm 0.064$ |
| ABIDE | UCLA_1 | TD | 27 | $13.52 \pm 1.95$ | 23 (85.2%) | $0.065 \pm 0.033$ |
| ABIDE | UCLA_2 | ASD | 8 | $12.35 \pm 2.06$ | 8 (100.0%) | $0.113 \pm 0.046$ |
| ABIDE | UCLA_2 | TD | 13 | $12.25 \pm 1.11$ | 11 (84.6%) | $0.095 \pm 0.065$ |
| ABIDE | UM_1 | ASD | 39 | $13.13 \pm 2.22$ | 31 (79.5%) | $0.109 \pm 0.071$ |
| ABIDE | UM_1 | TD | 46 | $13.63 \pm 2.93$ | 33 (71.7%) | $0.081 \pm 0.065$ |
| ABIDE | UM_2 | ASD | 13 | $14.88 \pm 1.55$ | 12 (92.3%) | $0.092 \pm 0.070$ |
| ABIDE | UM_2 | TD | 18 | $15.58 \pm 1.58$ | 17 (94.4%) | $0.059 \pm 0.030$ |
| ABIDE | USM | ASD | 16 | $16.23 \pm 1.86$ | 16 (100.0%) | $0.115 \pm 0.066$ |
| ABIDE | USM | TD | 13 | $13.82 \pm 2.49$ | 13 (100.0%) | $0.129 \pm 0.066$ |
| ABIDE | YALE | ASD | 25 | $12.91 \pm 3.11$ | 18 (72.0%) | $0.107 \pm 0.062$ |
| ABIDE | YALE | TD | 28 | $12.68 \pm 2.75$ | 20 (71.4%) | $0.092 \pm 0.059$ |
| ADHD200 | NYU | ADHD | 78 | $11.33 \pm 2.86$ | 64 (82.1%) | $0.102 \pm 0.044$ |
| ADHD200 | NYU | TD | 70 | $12.49 \pm 3.09$ | 39 (55.7%) | $0.096 \pm 0.044$ |
| ADHD200 | Peking | ADHD | 66 | $11.91 \pm 2.08$ | 61 (92.4%) | $0.127 \pm 0.039$ |
| ADHD200 | Peking | TD | 104 | $11.38 \pm 1.88$ | 70 (67.3%) | $0.122 \pm 0.049$ |
| HBN | HBNsiteCBIC | ASD | 5 | $10.42 \pm 1.89$ | 4 (80.0%) | $0.162 \pm 0.052$ |
| HBN | HBNsiteCBIC | ADHD | 147 | $11.94 \pm 3.14$ | 87 (59.2%) | $0.131 \pm 0.035$ |
| HBN | HBNsiteCBIC | TD | 23 | 11.72 ± 3.02 | 11 (47.8%) | 0.161 ± 0.049 |
| HBN | HBNsiteCUNY | ADHD | 7 | 10.92 ± 1.99 | 3 (42.9%) | 0.131 ± 0.011 |
| HBN | HBNsiteRU | ASD | 11 | 13.10 ± 3.18 | 7 (63.6%) | 0.142 ± 0.046 |
| HBN | HBNsiteRU | ADHD | 106 | 12.11 ± 2.99 | 72 (67.9%) | 0.133 ± 0.035 |
| HBN | HBNsiteRU | TD | 34 | 11.84 ± 2.90 | 21 (61.8%) | 0.147 ± 0.050 |
| HBN | HBNsiteSI | ASD | 7 | 12.39 ± 1.90 | 6 (85.7%) | 0.138 ± 0.037 |
| HBN | HBNsiteSI | ADHD | 44 | 12.81 ± 3.17 | 27 (61.4%) | 0.123 ± 0.038 |
| HBN | HBNsiteSI | TD | 32 | 11.71 ± 2.89 | 16 (50.0%) | 0.120 ± 0.048 |
| Global | All Sites | ASD | 325 | 12.90 ± 2.84 | 277 (85.2%) | 0.109 ± 0.063 |
| Global | All Sites | ADHD | 448 | 11.94 ± 2.92 | 314 (70.1%) | 0.125 ± 0.039 |
| Global | All Sites | TD | 613 | 12.47 ± 2.74 | 432 (70.5%) | 0.100 ± 0.056 |

**Notes:** FD, framewise displacement; ASD, autism spectrum disorder; ADHD, attention-deficit/hyperactivity disorder; TD, typically developing. The final three rows represent the consolidated demographic characteristics across all cohorts and sites following Combat harmonization protocols.

**Supplementary Table S2.**
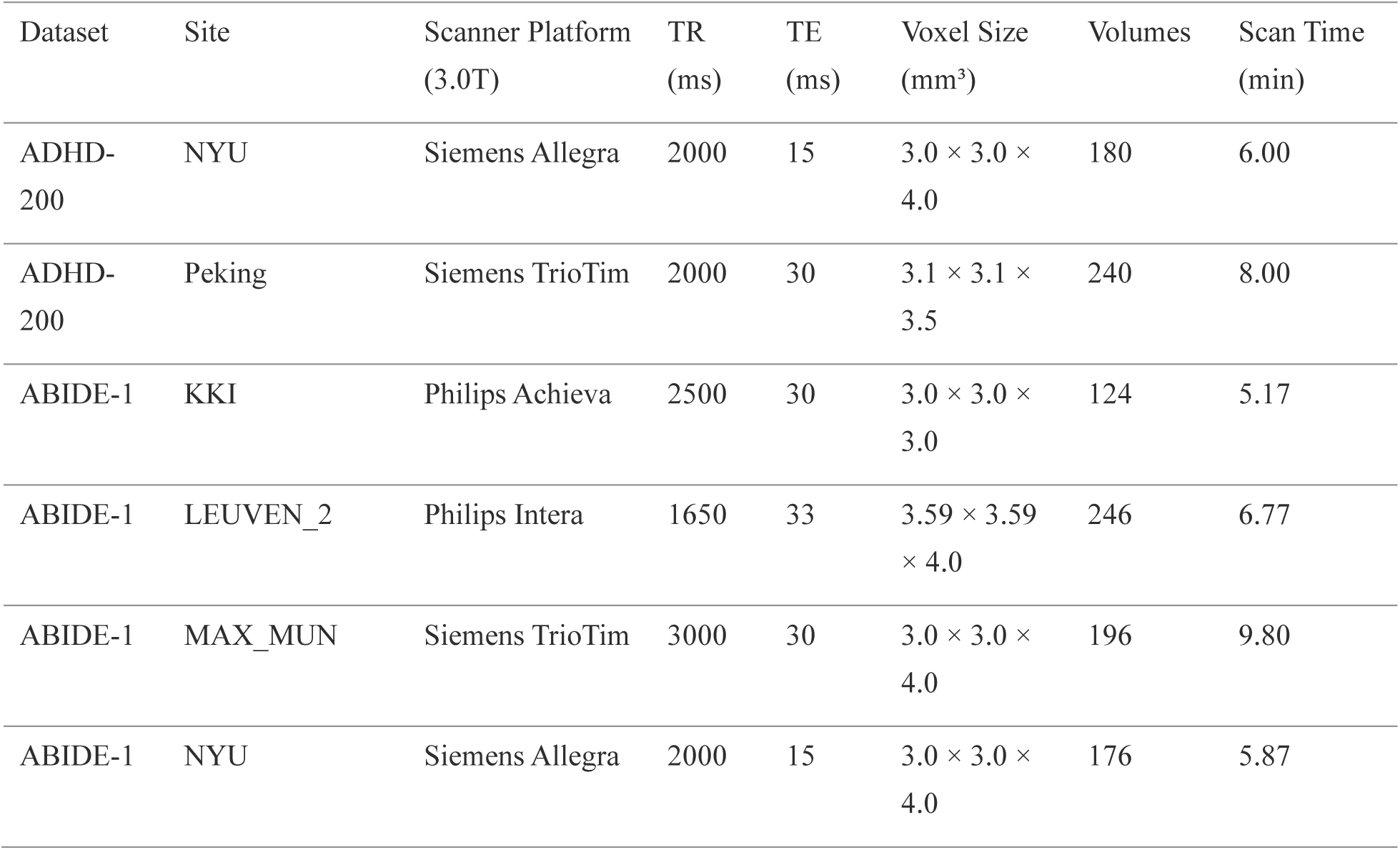
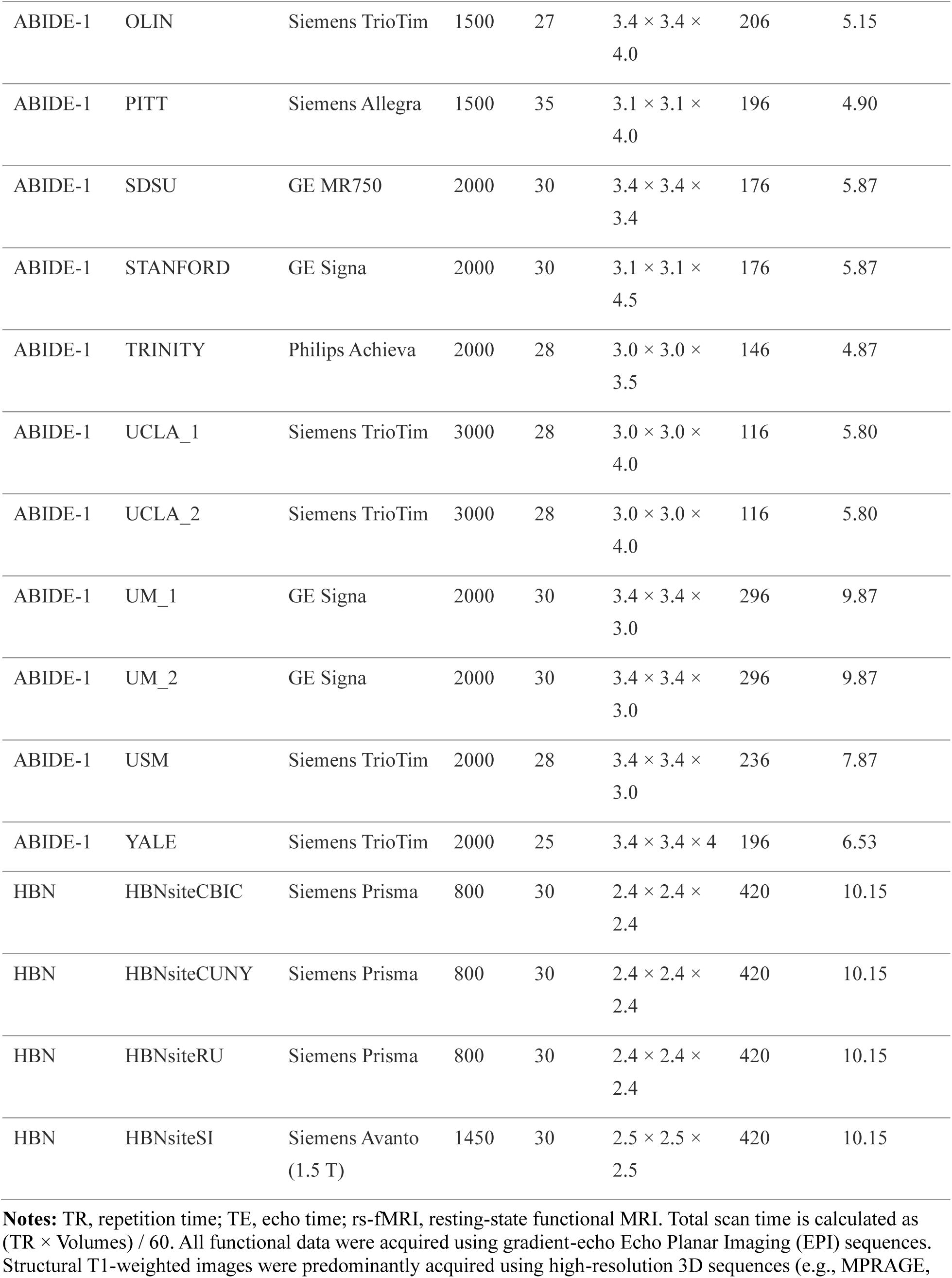
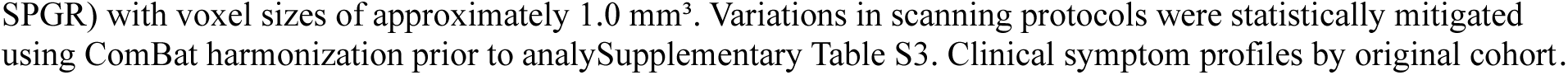
MRI acquisition parameters across independent scanning sites.

**Supplementary Table S3.** Clinical symptom profiles by original cohort.

| Cohort & Clinical Measure | ASD | ADHD | TD |
| --- | --- | --- | --- |
| ABIDE |  |  |  |
| ADOS Total Score | 12.06 ± 4.43 (n=254) | N/A | 1.38 ± 1.61 (n=13) |
| ADOS Social Affect | 9.06 ± 3.56 (n=235) | N/A | 1.00 ± 1.53 (n=13) |
| ADOS Restricted/Repetitive | 2.88 ± 1.77 (n=222) | N/A | 0.31 ± 0.85 (n=13) |
| ADHD-200 |  |  |  |
| ADHD Index | N/A | 60.95 ± 13.63 (n=144) | 36.00 ± 10.07 (n=160) |
| Inattentive | N/A | 50.98 ± 22.59 (n=144) | 28.13 ± 15.56 (n=160) |
| Hyper/Impulsive | N/A | 46.40 ± 24.93 (n=144) | 27.09 ± 16.39 (n=160) |
| HBN |  |  |  |
| SRS Total Score | 66.74 ± 11.19 (n=23) | 57.29 ± 10.65 (n=303) | 48.24 ± 8.00 (n=88) |
| SRS Social Affect | 192.30 ± 32.56 (n=23) | 168.46 ± 26.92 (n=303) | 147.10 ± 21.15 (n=88) |
| SRS Restricted/Repetitive | 64.74 ± 10.96 (n=23) | 56.23 ± 11.45 (n=303) | 47.17 ± 7.52 (n=88) |
| SWAN Total Score | -0.19 ± 1.04 (n=23) | 0.68 ± 0.84 (n=303) | -0.57 ± 1.05 (n=88) |
| SWAN Inattentive | -0.07 ± 1.17 (n=23) | 1.06 ± 0.94 (n=303) | -0.47 ± 1.18 (n=88) |
| SWAN Hyperactive | -0.31 ± 1.06 (n=23) | 0.30 ± 1.10 (n=303) | -0.67 ± 1.10 (n=88) |

**Notes:** Values are presented as Mean ± SD (number of valid observations). Symptom scores were acquired using cohort-specific psychometric instruments. Due to instrument heterogeneity and selective testing protocols (e.g., typically developing controls in ABIDE lacking ADOS scores), absolute scores cannot be aggregated across different cohorts. N/A indicates that the respective clinical measure was not collected for that diagnostic group within the specific cohort.

**Supplementary Table S4.**
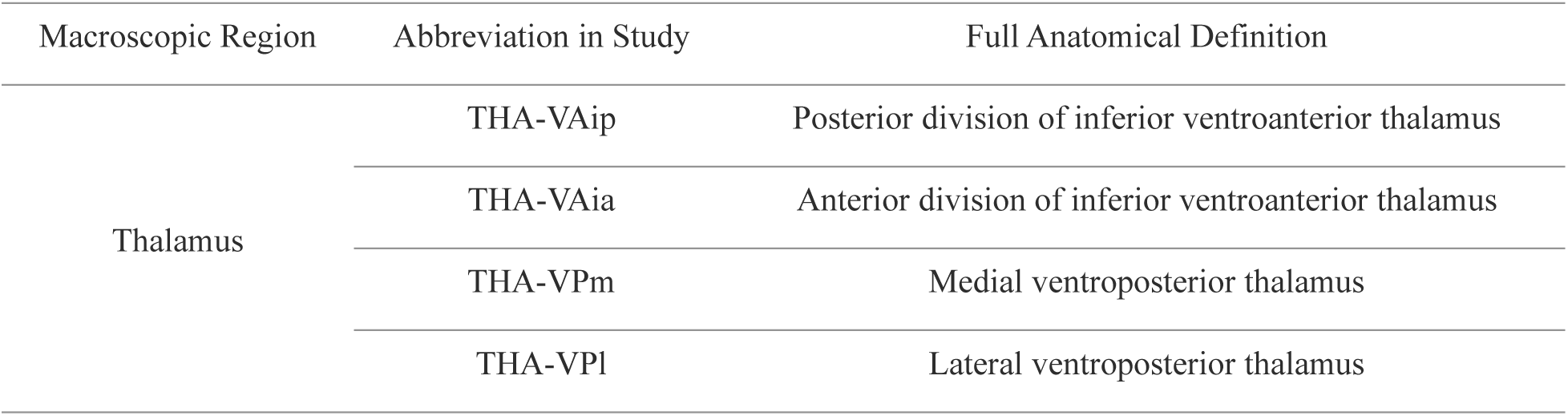
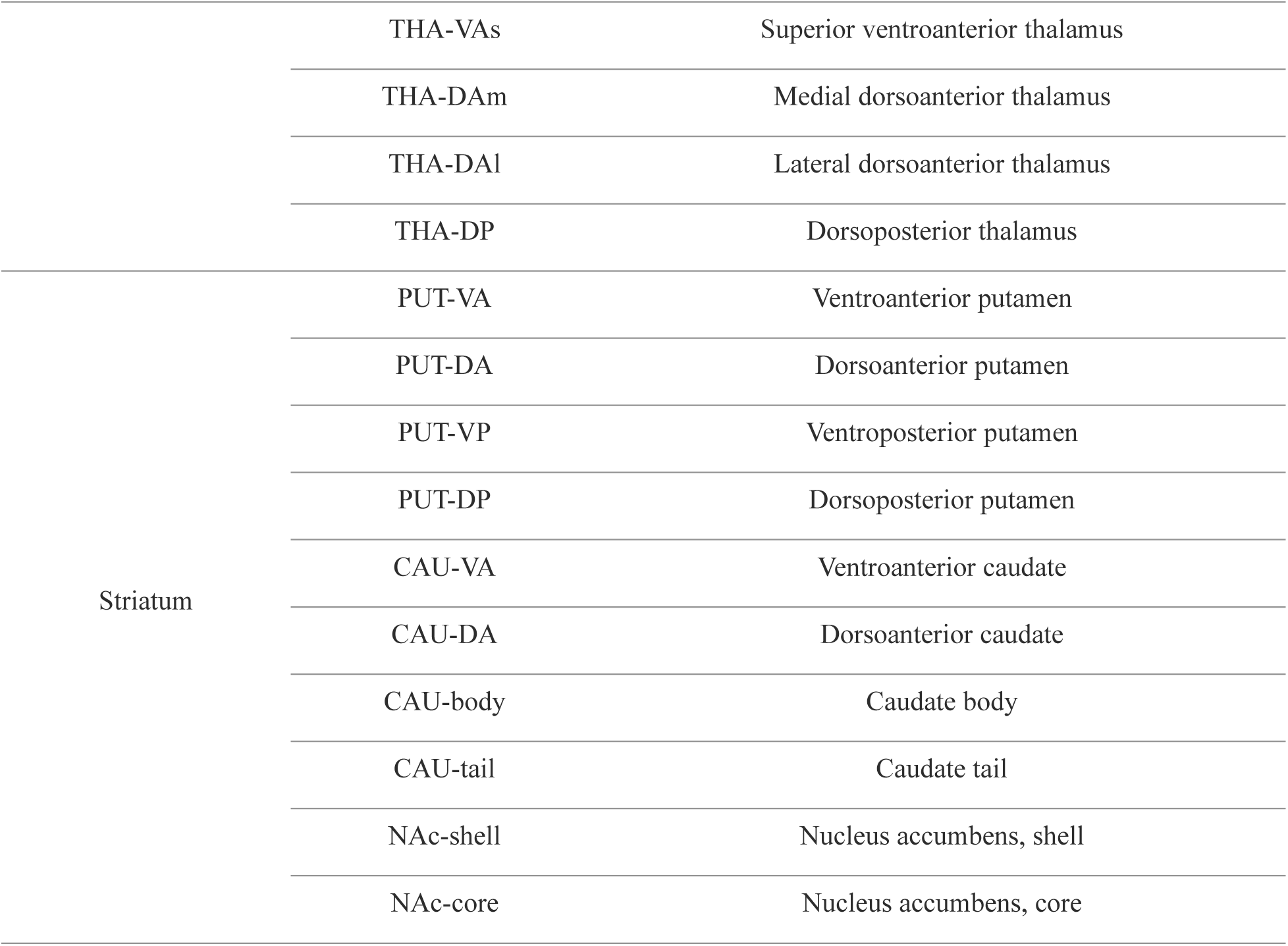
Nomenclature and Anatomical Definitions for Subcortical Subdivisions.

